# The sympatho-adrenal contribution to inferior vena cava formation

**DOI:** 10.1101/2025.08.01.668036

**Authors:** Aki Murashima, Daisuke Matsumaru, Takashi Moriguchi, Yuichi Shima, Miki Inoue, Chikako Yokoyama, Ken-ichirou Morohashi, Yoshiaki Kubota, Takatoshi Ueki, Jiro Hitomi, Sumio Isogai

## Abstract

Vascular development is fundamental during embryogenesis. Although the molecular and cellular mechanisms of vascular formation have been extensively studied, the developmental basis common to anatomical variations in major veins remains poorly understood^1–4^. In particular, the renal segment of the inferior vena cava (IVC) exhibits marked anatomical variability, and these structural differences often affect the safety of retroperitoneal surgeries^5,6^. Here we show that the renal segment of the IVC in mouse embryos does not arise from the subcardinal vein (SubCV), as classically proposed^7^, but instead forms from a vascular plexus in the para-aortic ridge (PAR) closely associated with developing sympathetic and adrenal tissues. Three-dimensional imaging with molecular markers revealed the vascular plexus in PAR arises independently of the SubCV and develops in Zuckerkandl organ and adrenal gland primordia. Moreover, genetic models demonstrated that the development of the sympathetic and adrenal tissues contributes to the organization and remodeling of this vascular network. This sympathoadrenal contribution contrasts with the classical mesonephros-centric model and explains the consistent IVC structures observed even in the absence of renal development. Our findings suggest a mammal-specific mode of IVC formation driven by the emergence of dominant sympathetic paraganglia and reduced reliance on mesonephric filtration. This revised developmental framework provides insight into the evolutionary divergence of venous anatomy and may help interpret clinically significant vascular variations during abdominal surgery.

## Main Text

Accumulating research on vascular formation has led to considerable progress in terms of molecular and cellular processes^1–4^. However, the mechanisms underlying vascular patterning with broad morphological variations have been poorly elucidated. Particularly, the developmental processes of the renal segment of the inferior vena cava (IVC), which encompasses the renal, adrenal, gonadal, and retroperitoneal veins, remain unclear. Extensive anatomical variations of the renal segment of the IVC around the abdominal aorta often lead to life-threatening hemorrhage during a wide range of retroperitoneal surgeries including nephrectomy, kidney transplantation, abdominal aortic aneurysm repair, and para-aortic lymphadenectomy^5,6^. Elucidating the developmental mechanisms underlying these variations in the renal segment of the IVC will advance our understanding of embryonic venous patterning and ultimately contribute to safer surgical approaches to the retroperitoneal region.

The development of the IVC is explained by the anastomosis and selective development of the bilateral primitive venous systems—the posterior cardinal vein (PostCV), subcardinal vein (SubCV), and supracardinal vein (SupraCV)—along the trunk, regulated by blood flow dynamics^7^. The SubCV forms in close association with the mesonephros, excretory primordium transiently formed in mid-gestational embryos^7–9^. As the fetal mesonephros develops and functions, inter-SubCV anastomosis is believed to form the renal segment of the IVC branching the renal, adrenal, and gonadal veins. In classical reevaluations, the ambiguity of SubCV has often been described in mammals, where the development of the mesonephros is not prominent^7,10^. Furthermore, in humans, even when the left renal vein is absent due to abnormal renal development, the renal segment of the IVC receiving the left suprarenal and inferior phrenic veins is consistently observed. However, the SubCV model does not fully explain these observations.

### SubCV formation in murine embryos

The developmental processes of the venous system have been well documented using classical three-dimensional (3D) reconstruction, ink injection, and corrosion casting^7,8,10–14^. Given this, we center on the vascular plexus identified by Endomucin (EMCN), an endothelial cell marker that is predominantly expressed in the venous and premature plexuses, which enables the analysis of the vascular plexus regardless of the presence or absence of a lumen^15^.

PostCV formed dorsally to the mesonephros along the cranio-caudal axis at E11.5 (Fig. 1A). The SubCV (longitudinal anastomosis of the mesonephric venous sinuses surrounding the mesonephric tubules) was only present in the cranio-medial region of the mesonephros (Fig. 1A, yellow arrowheads). At the level caudal to the superior mesenteric artery (SMA), condensed mesenchyme was observed without differentiation of the mesonephric tubules (Fig. 1A, asterisks). SubCV did not develop in this region, leaving discontinuous and lumenless endothelial cell distribution (Fig. 1A, white arrows). The endothelial cells located ventral to the aorta formed scattered fine tributaries emerging from the mesenteric root, connecting with the SubCV or PostCV above and below the origin of the SMA; however, the tributaries did not form an inter-SubCV vascular network (Fig. 1A, yellow arrows). Cranial SubCVs were separated from the mesonephroi by the body cavity at E12.0. On the right side, it connected with the hepatic segment of the “posterior” (equivalent to the “inferior” in the case of humans) vena cava (PVC_hep_), while on the left side, it anastomosed with the PostCV (Fig. 1B). PostCVs gave rise to numerous ventral branches, forming a vascular plexus in the para-aortic ridge (PAR), including the ventral side of the aorta from the SMA to its caudal region (Fig. 1B, white arrowhead). At E12.5, the vascular plexus developed further and the gonadal venous primordium draining from the caudal pole of the gonad to the caudal edge of this network became visible. Additionally, branches returning from the peri-metanephric vascular network and caudal mesonephric duct drained into this plexus (Fig. 1C). At E13.5, a dominant vessel emerged within the vascular plexus, forming a conduit that linked the left-sided venous drainage to the PVC_hep_. As the metanephroi ascended and their hilum adducted, their drainage pathways (the renal veins), which initially connected dorsally to the left and right gonadal veins, shifted to connect from the lateral side (Fig. 1C and D, broken lines). The gonadal and renal veins formed common trunks which then connected to the conduit, forming the anlage of the definitive renal vein. On the left side, the plexus also received the inferior phrenic vein in its cranial plane. Consequently, branches of the future definitive PVC renal segment were constructed at this stage (Fig. 1D, broken lines). These results indicate that a vascular plexus formed in the PAR, and not in the site of inter-SubCV anastomosis, thus contributing to the development of PVC in the renal segment.

**Fig. 1.**
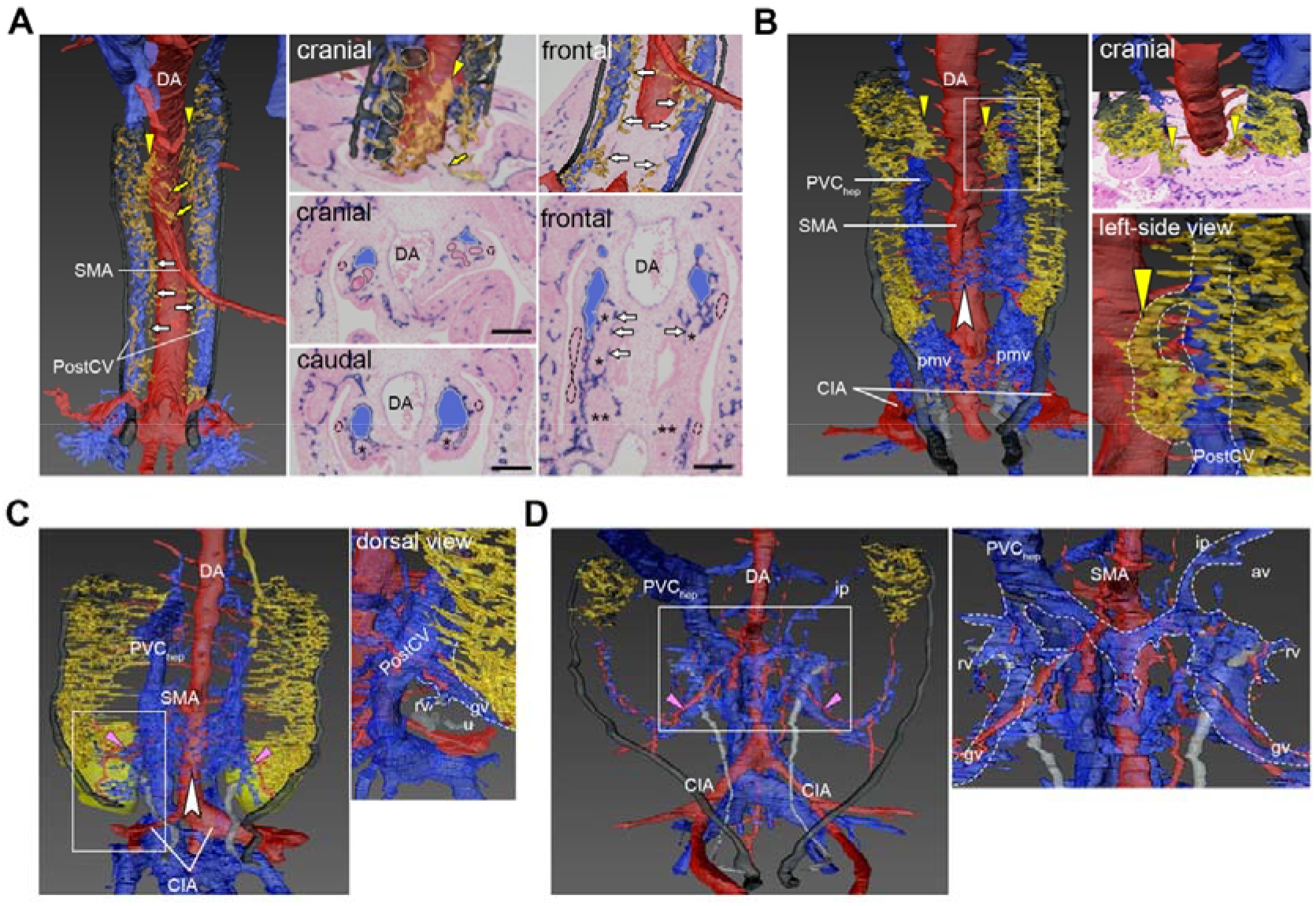
Three-dimensional reconstruction for EMCN-stained vascular systems based on serial sections at E11.5 (A), 12.0 (B), 12.5 (C) and 13.5 (D). Arterial (red) and venous (blue) system, mesonephric sinusoid (yellowish brown), nephric duct/tubules (black) and ureter (gray) were reconstructed. Right-side and frontal view of the reconstructed cranial mesonephros with the original slice (A and B). The enclosed areas in B, C and D are enlarged. The veins outlined by broken lines in D represent the branches of the future definitive PVC renal segment. Yellow arrowheads, SubCV; yellow arrow, endothelial cells in mesenteric root; white arrowhead, the vascular plexus in the PAR; pink arrowhead, the gonadal artery; white arrow, rudimental endothelial alignment in caudal mesonephros; black asterisk, mesenchymal condensation (caudal mesonephros); double asterisk, mesenchymal condensation (metanephros); dotted line, the nephric duct/tubule; SMA, superior mesenteric artery; CIA, common iliac artery; DA, dorsal aorta; PVC_hep_, hepatic segment of posterior vena cava; gv, gonadal vein; rv, renal vein; ip, inferior phrenic vein; av, adrenal vein; pmv, peri-metanephric vascular network; k, kidney; Scale bars indicate 100 µm.

### Vascular plexus formation in the para-aortic ridge

We previously reported that definitive renal arteries develop in close association with sympathetic paraganglia, such as the adrenal medulla and the Zuckerkandl organ (ZO), in PAR^13^. To test the developmental relationship between the sympathetic tissues and PVC, vascular formation in the PAR was analyzed. The sympathetic tissues are marked by Tyrosine hydroxylase (TH) immunoreactivity during its development, and Cocaine- and amphetamine-regulated transcript (CART) discriminates the sympathetic ganglia from the paraganglia^18^. TH-positive cells were observed in the sympathetic chain, and they expanded to the ventral PAR surrounding the caudal half of the SMA at E11.5, differentiating into ZO (Fig. 2A and B). Several arteries directed towards the mesonephros were observed bilaterally along the ventral wall of the dorsal aorta at E11.5, gradually decreasing in diameter (Fig. 2C, orange arrowheads). As the vascular plexus in the ZO became evident, the development of the gonadal arteries became more prominent than that of the mesonephric arteries, supplying the ZO through its tributaries (Fig. 2C, white arrowheads). At this stage, a prominent vascular network developed on the cranial and ventral surfaces of the TH-high bilateral ZO but not in the TH-low and CART-high prevertebral mesenteric ganglia (Fig. 2A and D black arrowheads, and E). Cell lineage analyses using *Cdh5-Cre*^*ERT2*^;*ROSA-tdTomato* mice showed that the vascular network in PAR was composed almost entirely of endothelia differentiated prior to plexus formation, suggesting a developmental mechanism of angiogenesis from PostCV (Fig. 2F).

**Fig. 2.**
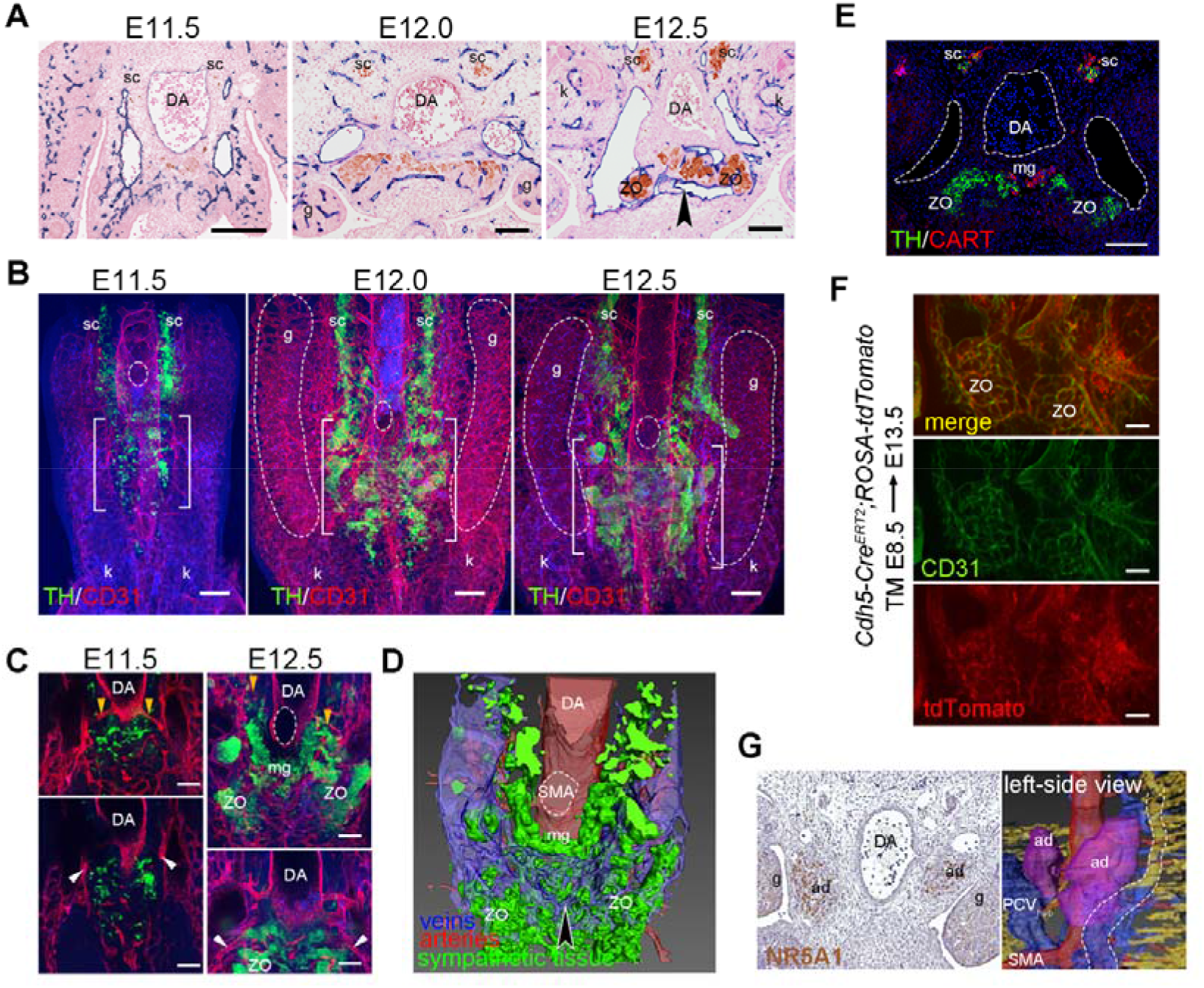
The para-aortic ridge (PAR) and vascular development. (A)Immunohistochemical staining of TH indicating sympathetic neuroblast and ZO was co-stained with EMCN. (B)Whole-mount immunofluorescence of vascular structure (CD31, red) with the sympathetic tissue (TH, green). The white bracket shows the ZO developing ventral to the aorta. (C) Mesonephric (orange arrowheads) and gonadal (white arrowheads) arteries in B are enlarged. (D) Three-dimensional reconstruction of the E12.5 specimen in E. (E) Immunofluorescence showing ZO (CART-negative/TH-high) and mesenteric ganglia (CART-high/TH-low). (F) Immunofluorescent staining for CD31 in *Cdh5-Cr*^*eERT2*^;*ROSA-tdTomato* embryo at E13.5; TM was given at E8.5. (G) Immunohistochemically for NR5A1 at E12.5. Left-side view of the reconstructed adrenal primordia located between the SubCV (behind the ad, see also the Supplemental Movie E12.5) and PVC_hep_ on the right side, and the SubCV (broken line, yellowish brown) and PostCV (broken line, blue) on the left side. DA, dorsal aorta; SMA, superior mesenteric artery; k, kidney; g, gonad; sc, sympathetic chain; mg, mesenteric ganglia; ad, adrenal primordia. Scale bars indicate 100 μm (A, C, E, F) and 200 µm (B).

Adrenal primordia develop bilaterally on the cranial side of the ZO^19^. The adrenal cortex labeled by NR5A1 was located at the site of anastomosis between the SubCV and PVC_hep_ on the right side or PostCV on the left side, with its returning pathway draining cranially into the vascular network of the ZO (Fig. 2G).

### Experimental modification of nephric and PAR development

To experimentally verify the contribution of the nephric and/or sympathoadrenal organs to PVC development, we conducted analyses using genetically modified mice. *Hoxb7-Cre;ROSA-DTA* mice exhibit severe defects in the urogenital system, including the metanephros, due to cell death in the nephric duct (ND), particularly in the caudal region of the mesonephros^20^. In this mutant, the vascular plexus in the PAR formed as observed in the control, and the conduit was established as a drainage pathway for the left gonad and adrenal gland despite the retardation in meso- and metanephric development (Fig. 3A). *Gata3* knockout (KO) mice exhibit defects in the mesonephros and metanephros along with impairments in the differentiation of TH-positive sympathetic neurons and chromaffin cells^21,22^. We found that the vascular plexus in the ZO was poorly developed in *Gata3* KO embryos relative to littermate controls. The conduit was formed dorsally to the ZO region, with a delayed onset and more cranial displacement than the controls at E14.5 (Fig. 3B). To exclude the effects of meso- and metanephric defects in *Gata3* KO mice, we used *P0-Cre* mice^23^ to generate neural crest cell-specific *Gata3* (*Gata3*^*NC*^) KO mice and assessed vascular plexus development. The plexus development was sparse in the mutants in comparison to controls at E13.5, and the conduit was formed dorsal to the ZO at E14.5, as observed in the *Gata3* conventional KO embryos (Fig. 3B and C). Additional KO of *Nr5a1* (*Nr5a1;Gata3*^*NC*^) for adrenocortical hypoplasia^24^ firmly represented the altered venous orbit (Fig. 3D). These findings suggest that the midline vascular plexus, which contributes to PVC formation, is influenced by the development of ZO and sympatho-adrenal tissues in the PAR.

**Fig. 3.**
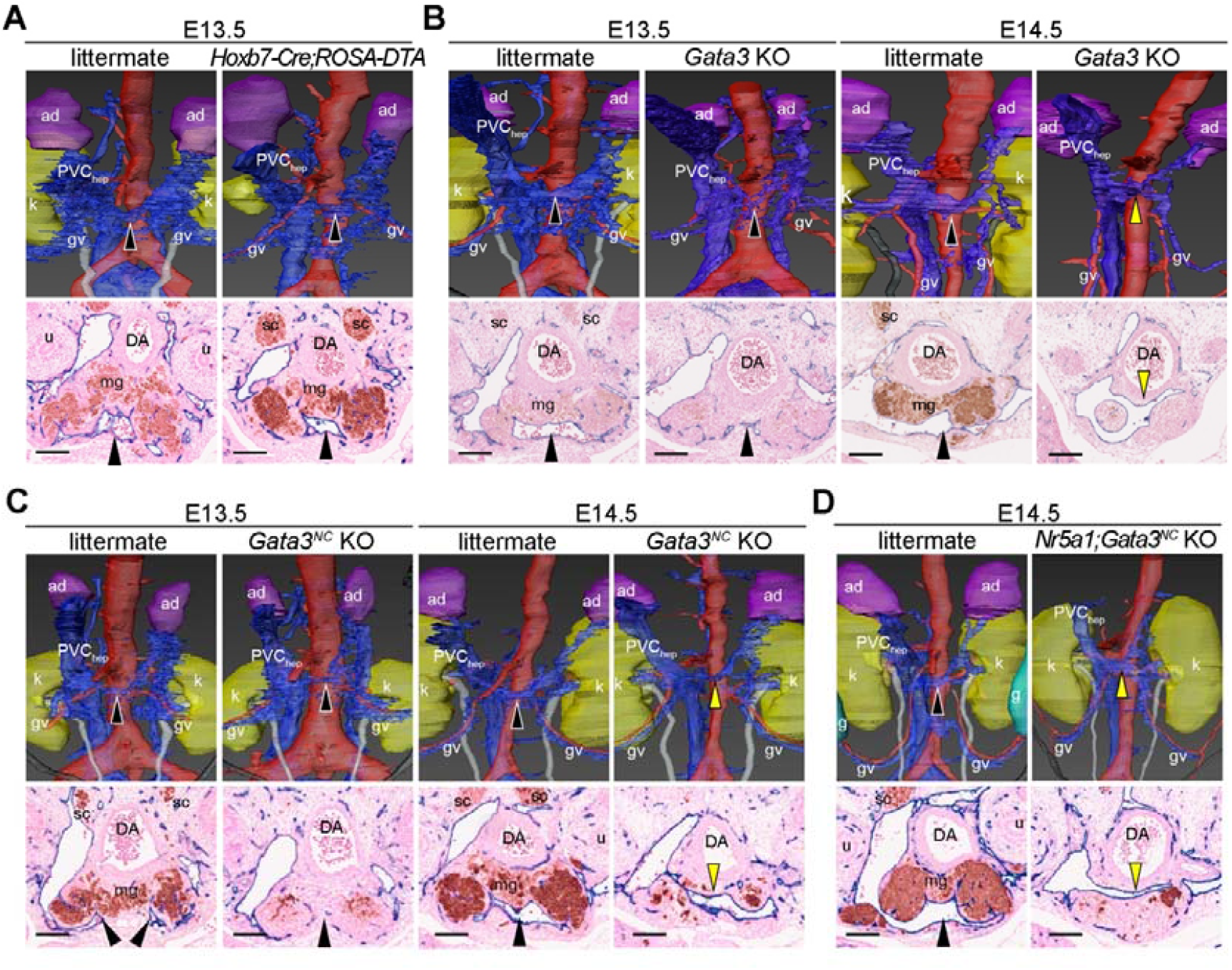
Sympathoadrenal organ development in the PAR affects the formation of the renal segment of the PCV. The developing venous systems of *Hoxb7-Cre;ROSA-DTA* (A), *Gata3* KO (B), *Gata3*^*NC*^ KO (C) and *Nr5a1;Gata3*^*NC*^ KO (D) embryos at E13.5 and E14.5 were immunohistochemically stained with anti-EMCN (blue) and anti-TH (brown) antibodies, and serial sections were reconstructed three dimensionally. Representative slices showing ventral (black arrowhead) and dorsal (yellow arrowhead) conduits in ZO are indicated. DA, dorsal aorta; SMA, superior mesenteric artery; ip, inferior phrenic vein; gv, gonadal vein; k, kidney; g, gonad; mg, mesenteric ganglia; ad, adrenal primordia. Scale bar: 100 µm.

## Discussion

The conventional model suggests that the renal segment of the IVC originates from the SubCV, which drains the mesonephros (Fig. 4A). This model has been substantiated in amphibians, where the mesonephros develops into a permanent kidney, and in birds and reptiles, where its transient development is more pronounced than in mammals^25–28^. In this study, we demonstrated that the SubCV is highly underdeveloped in mouse embryos, which is occasionally noted in human embryos^10^. In mouse embryos and other mammals including humans, placental development reduces reliance on embryonic kidneys for blood filtration, rendering the formation of a renal portal system unnecessary.

**Fig. 4.**
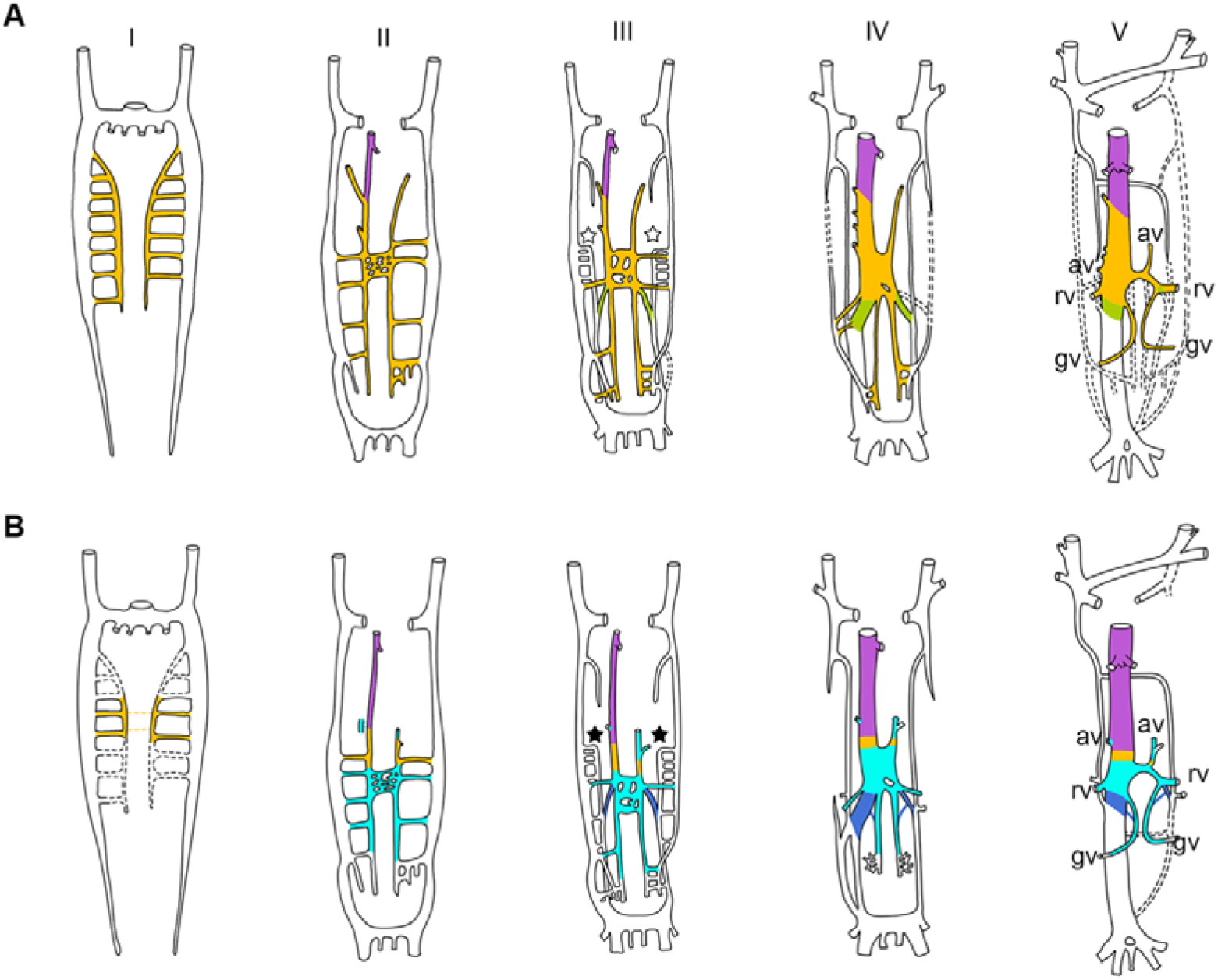
Standard and novel models of inferior vena cava development. (A) The standard model illustrating the venous system development in human embryos at 4 mm (I), 11 mm (II), 15 mm (III), and 16 mm (IV) stages, as well as in the adult (V) adapted from McClure and Butler^7^. In this model, the renal segment of the IVC originates from the mesonephric venous system (yellow). Paired longitudinal veins ⍰ represent the SupraCV, and light green veins indicate anastomoses between the SubCV and SupraCV. (E) The novel model representing an alternative origin of the renal segment from the para-aortic venous system (cyan). The veins ⍰ correspond to veins accompanying the sympathetic trunks. The blue veins indicate anastomoses between the para-aortic venous system and the infrarenal segment of the IVC. Abbreviations as in Fig. 1.

The 3D imaging and molecular marker-based approaches used in this study enabled precise visualization of prevertebral sympathetic tissues and venous structures. Considering the distinct developmental trajectories across vertebrates, we propose that the development of the prevertebral sympathetic tissues and adrenal gland in the PAR plausibly integrates bilateral venous return from the trunk (Fig. 3E). The prominent sympathetic paraganglia composed of aggregated chromaffin cells, ZO, have been reported mainly in mammalian embryos^29,30^, whereas in other vertebrates, chromaffin cells are broadly distributed around the aorta and are either isolated or clusters of a few cells^31^. The decline in mesonephric development and the subsequent acquisition of predominant sympathetic paraganglia in mammals may play a pivotal role in the development of the vena cava in vertebrate phylogeny. Our model provides a neurovascular perspective on IVC formation, offering a new framework for interpreting anatomical variations that cannot be fully explained by traditional mesonephric-derived models.

## Materials and Methods

### Mice

*Gata3-LacZ*^27^ (kindly provided by Dr. Engel), *Gata3-flox*^28^, *Nr5a1*^21^ and *Cdh5-Cre*^*ERT2* 29^ mice used in this study have been described previously. *P0-Cre*^20^ mice were obtained from the CARD mouse bank at Kumamoto University. *Hoxb7-Cre*^30^, *ROSA-DTA*^31^ and *ROSA-tdTomato*^32^ were obtained from the Jackson Laboratory (stock 004692, 010527, and 007914). ICR mice were obtained from Japan SLC, Inc. (Hamamatsu, Japan). Mice were maintained under normal light/dark conditions. Noon on the day on which the vaginal plug appeared was designated as embryonic day (E) 0.5. Embryos were collected between E11.5 and E14.5, and fixed overnight in 4% paraformaldehyde dissolved in phosphate-buffered saline (PBS). Pharmacological rescue of *Gata3* mutant embryos has been previously described^27^. Tamoxifen (TM, Sigma, St. Louis, MO, USA) was dissolved in sesame oil (Kanto Chemical, Tokyo, Japan) at a final concentration of 10 mg/ml. Four milligrams of TM per 40 g body weight was administered (ip) to pregnant mice^33^. Experiments using laboratory mice were approved by the Committee on Animal Research at Iwate Medical University. Embryos and pups for each experiment were collected from more than three independent pregnant females.

### Immunohistochemistry for serial paraffin sections

Embryos were dehydrated and embedded in paraffin. Serial sections through the body trunk were prepared at a thickness of 6–7 µm. After deparaffinization and hydration, the sections were washed with distilled water and autoclaved at 121°C for 1 min in citrate buffer for antigen retrieval. Endogenous peroxidase activity was quenched with 3% H_2_O_2_. Non-specific antibody binding was blocked by pre-incubating the specimens in 1% FBS/PBS. The sections were stained for 1 hour with primary antibodies at room temperature. The primary antibodies were detected by HRP- or AP-conjugated secondary antibodies, followed by incubation with 3,3’-diaminobenzidine (DAB) (040–27001; Wako, Osaka, Japan) and NBT/BCIP stock solution (Roche, Basel, Switzerland) for 10–30 min. The sections were washed and subjected to nuclear staining with Nuclear Fast Red (ScyTek Laboratories, Inc., UT, USA). Subsequently, the sections were dehydrated, permeated and mounted using Malinol (Muto Pure Chemicals, Tokyo, Japan). As a negative control, the specimens were stained without the primary antibodies (data not shown).

### Whole-mount immunofluorescence

Embryos were washed three times in PBSTx (0.1% TritonX-100 in PBS), blocked with 10% FBS/PBSTx for 90 min, and then incubated with the primary antibodies diluted in the blocking solution for three days. Embryos were washed in PBSTx, incubated for three days with secondary antibodies, and washed in PBSTx. Specimens were then stained with Hoechst 33342 (Sigma-Aldrich, Osaka, Japan) for 1 hour, soaked in 50% Tissue-Cleaning Reagent CUBIC-R+ [for animals] (CUBIC-R+, T3741; Tokyo Chemical Industry, Tokyo, Japan) overnight, and replaced with 100% CUBIC-R+. A fluorescence microscope (BZ800, Keyence, Osaka, Japan) or a confocal laser scanning microscope (A1R, Nikon, Tokyo, Japan) was used to obtain Z-stack images.

The primary antibodies used in this study were anti-EMCN (sc-65495, Santa Cruz Biotechnology, CA, USA), anti-CD31 (AF3628, R&D Systems, MN, USA), anti-TH (AB152, MAB138, Merck Millipore, MT, USA), anti-CART (H-003-62, Phoenix Pharmaceuticals, CA, USA), and anti-NR5A1^34^ The secondary antibodies used were anti-rat AP (15-16-06, LGC Clinical Diagnostics, MD, USA), anti-rabbit HRP (ab6721, Abcam, Cambridge, UK), anti-rabbit Alexa546, anti-rabbit Alexa488, anti-mouse Alexa488, anti-mouse Alexa546 and anti-goat Alexa647 (A11010, A11008, A21206, A11001, A11003, A21447, Thermo Fisher Scientific, MA, USA).

### Three-dimensional reconstruction

Serial sections were scanned using a digital slide scanner (Nanozoomer S210, Hamamatsu Photonics, Japan) and exported serial images were converted into an image sequence file (ImageJ, NIH, USA). The files were processed using the Amira software program (version 5.6, Maxnet Co, Ltd, Tokyo, Japan). Images were aligned and segmented, followed by the manual delineation of the blood vessels and structures of interest. All reconstructed images are shown in the Supplemental Movies (Movie S1-S4).

## Acknowledgments

We thank Satoko Kumagai, Michiyo Tashiro, Kinji Ishida and staff in the Technical Support Center for Life Science Research, Iwate Medical University for the technical assistances. We also appreciate Drs. Douglas Engel (University of Michigan Medical School) and Dr. Sho Ohta (National Institutes of Health) for resource and advise on 3D reconstruction techniques. We use confocal microscopy and Digital slide scanners in Technical Support Center for Life Science Research, Iwate Medical University and Nagoya City University. This study was supported by JSPS KAKENHI Grant Number 20K07229, 17K08495 and KEIRYOKAI Research Grant 135.

## Author contributions

Conceptualization: A.M., S.I.; Data acquisition: A.M, D.M., M.I.; Data analysis: A.M., S.I; Resources: T.M., M.I., K.M., Y.S., J.H., T.U.; Funding acquisition: A.M., J.H.; Writing - original draft: A.M.; Writing - review & editing: all authors.

## List of Supplementary Materials

Movies S1 to S4

## Movie captions

Movie S1.

Three-dimensional reconstruction for EMCN-stained vascular systems based on serial sections of mouse embryos at E11.5.

Movie S2.

Three-dimensional reconstruction for EMCN-stained vascular systems based on serial sections of mouse embryos at E12.0.

Movie S3.

Three-dimensional reconstruction for EMCN-stained vascular systems based on serial sections of mouse embryos at E12.5.

Movie S4.

Three-dimensional reconstruction for EMCN-stained vascular systems based on serial sections of mouse embryos at E13.5.

